# Extracellular Vesicle Antibody Microarray for Multiplexed Inner and Outer Protein Analysis

**DOI:** 10.1101/2022.04.05.487234

**Authors:** Rosalie Martel, Molly L. Shen, Philippe DeCorwin-Martin, Lorenna Oliveira Fernandes de Araujo, David Juncker

## Abstract

Proteins are found both outside and inside of extracellular vesicles (EVs) and govern the properties and functions of EVs, while also constituting a signature of the cell of origin and of biological function and disease. Outer proteins on EVs can be directly bound by antibodies to either enrich EVs, or probe the expression of a protein on EVs, including in a combinatorial manner. However, co-profiling of inner proteins remains challenging. Here, we present the high-throughput, multiplexed analysis of extracellular vesicle inner and outer proteins (EVPio). We describe the optimization of fixation and heat-induced protein epitope retrieval for EVs, along with oligo-barcoded antibodies and branched DNA signal amplification for sensitive, multiplexed and high-throughput assays. We captured 4 subpopulations of EVs from colorectal cancer cell lines HT29 and SW403 based on EpCAM, CD9, CD63 and CD81 expression, and quantified the co-expression of 8 outer (integrins and tetraspanins) and 4 inner (heat shock, endosomal and inner leaflet) proteins. The differences in co-expression patterns were consistent with the literature and known biological function. In conclusion, EVPio analysis can simultaneously detect multiple inner and outer proteins in EVs immobilized on a surface, opening the way to extensive combinatorial protein profiles for both discovery and clinical translation.

---

Extracellular vesicles (EVs) are bound by a lipid bilayer, range from ∼50 nm to a few micrometers in size, and are released by cells in their environment. They include microvesicles, resulting from the budding of the cell plasma membrane; exosomes, small vesicles formed through the inner budding of endosomes which then fuse with the cell membrane; and apoptotic vesicles, shed during programmed cell death.^1, 2^ They carry a complex biomolecular cargo of lipids, nucleic acids and proteins, and have become recognized for their major role in cell-to-cell communication^3-7^ and as cellular messengers transferring material and molecular information from cell to cell. EVs are critical to many physiological processes and diseases, such as cancer,^8-11^ and have further been recognized for their role as biomarkers, notably based on variations of their protein content.^12-14^ There has been a growing interest in understanding which proteins are expressed and enriched in EVs, and which ones are co-enriched.

Various EV protein analysis technologies have been developed or optimized in recent years, notably mass spectrometry,^15, 16^ flow cytometry,^17, 18^ microfluidics,^19, 20^ plasmonics,^21, 22^ and antibody microarrays (as well as affinity binder arrays).^23, 24^ Many analyses rely on the bulk lysis of the EVs and detect both intravesicular proteins in the lumen of the vesicle (inner)^16, 25^ and membrane proteins that are accessible from the outside (outer). However, co-expression information is lost as part of the lysis and only ensemble averages are available. Enrichment by affinity capture of an outer protein followed by elution and/or lysis reveals the co-expression and enrichment for one outer protein at a time.^23, 26, 27^

A variety of proteins from multiple families have been probed in EVs. Tetraspanins and integrins are among the most commonly detected outer targets,^28^ while inner proteins such as cytokines,^29^ pathway effectors and cell machinery components were analyzed. Heat shock proteins, such as HSP90 and HSP70, and Alix, a protein involved in the syndecan-synthenin-Alix pathway for exosome biogenesis and cargo-sorting, are commonly found in EVs.^30, 31^

Antibody microarrays can be used for the pairwise combinatorial analysis of protein co-expression in EVs. Arrays of antibody microspots targeting outer EV proteins (*capture antibodies*) are used to enrich EVs, forming spatially resolvable subpopulations based on outer protein expression. Immobilized EVs can then be probed with additional, labeled *detection antibodies* against other transmembrane and surface proteins. However, the co-expression of outer-inner proteins has remained challenging because the bilayer shields the inner proteins which are not accessible for antibody binding.

By contrast, in tissues and cells, extracellular and cytosolic proteins are readily mapped using immunohistochemistry and immunocytochemistry, respectively. To stain cytosolic proteins, including the cytosolic epitopes of transmembrane proteins, cells are chemically fixed (cross-linked) and permeabilized using surfactants, followed by a so-called antigen retrieval (AR) step that breaks cross-links and exposes protein epitopes for binding, using enzymes, chemicals such as urea, heat treatments, or a combination thereof.^32-34^ Conceptually, it may be possible to apply similar protocols to EVs. To the best of our knowledge, this had not yet been reported, possibly because EVs are much smaller, more susceptible to disruption, and contain comparatively little protein. Therefore, a distinct protocol specific to EVs is needed.

Here, we introduce the high-throughput, multiplexed analysis of extracellular vesicle inner and outer proteins (EVPio). Using antibody microarrays, (i) EV subpopulations are captured and immobilized based on the expression of an outer target protein, (ii) fixed and subjected to AR using protocols developed and optimized here, followed by (iii) the multiplexed detection of up to three inner and outer proteins per spot, measured simultaneously using different fluorophores. The signal-to-noise ratio (SNR) was derived by dividing the assay signal by the background noise (embodied by the standard deviation, SD, of replicate measurements) and used to benchmark and optimize most assay steps by maximizing SNR. To address the low concentration and weak signal of inner proteins, we introduce DNA barcoding of the detection antibodies and complementary, fluorescent, branched DNA structures for signal amplification. To illustrate the phenotyping potential of EVPio, EVs from two colorectal cancer (CRC) epithelial adenocarcinoma cell lines with distinct mutation profiles, HT29 and SW403,^35^ were captured using four outer proteins, and the pairwise co-expression of 8 outer and 4 inner proteins quantified in each of the four subpopulations.

## Results and Discussion

### EVPio Analysis: Overview and Operation Workflow

We introduce the combinatorial analysis of extracellular vesicle inner and outer proteins as a method to study the inner and outer protein contents of EVs. EV subpopulations are enriched based on their surface protein content using antibody microarrays with multiple different capture antibodies (Figure 1). Briefly, using an inkjet spotter, each slide is patterned with 16 identical microarrays with each 100 spots of up to 10 different antibodies targeting surface protein markers, laid out as rows of 10 replicate spots. The microarrays were delimited by gaskets to form wells, and each well was incubated overnight with 50 μL of sample containing 1 × 10^10^ EVs/mL (obtained by size exclusion chromatography of cell supernatant). The EVs were fixed, then subjected to an AR step described in detail below, optimized to allow the simultaneous detection of multiple inner and outer proteins. Processed EVs were incubated with a mixture of up to three different detection antibodies per well, and upon rinsing and drying, binding of the detection antibodies was detected using a microarray scanner used to analyze the different fluorescence wavelengths. The overall protocol, from array fabrication to imaging, could be completed in three days. Many inner proteins are challenging to detect due to low concentrations and resulting weak signals. The detection antibodies were conjugated with an oligonucleotide barcode for hybridizing complementary fluorescent detection oligos, including linear and multi-branched oligos for further increasing the SNR,^36-38^ as detailed below.

**Figure 1.**
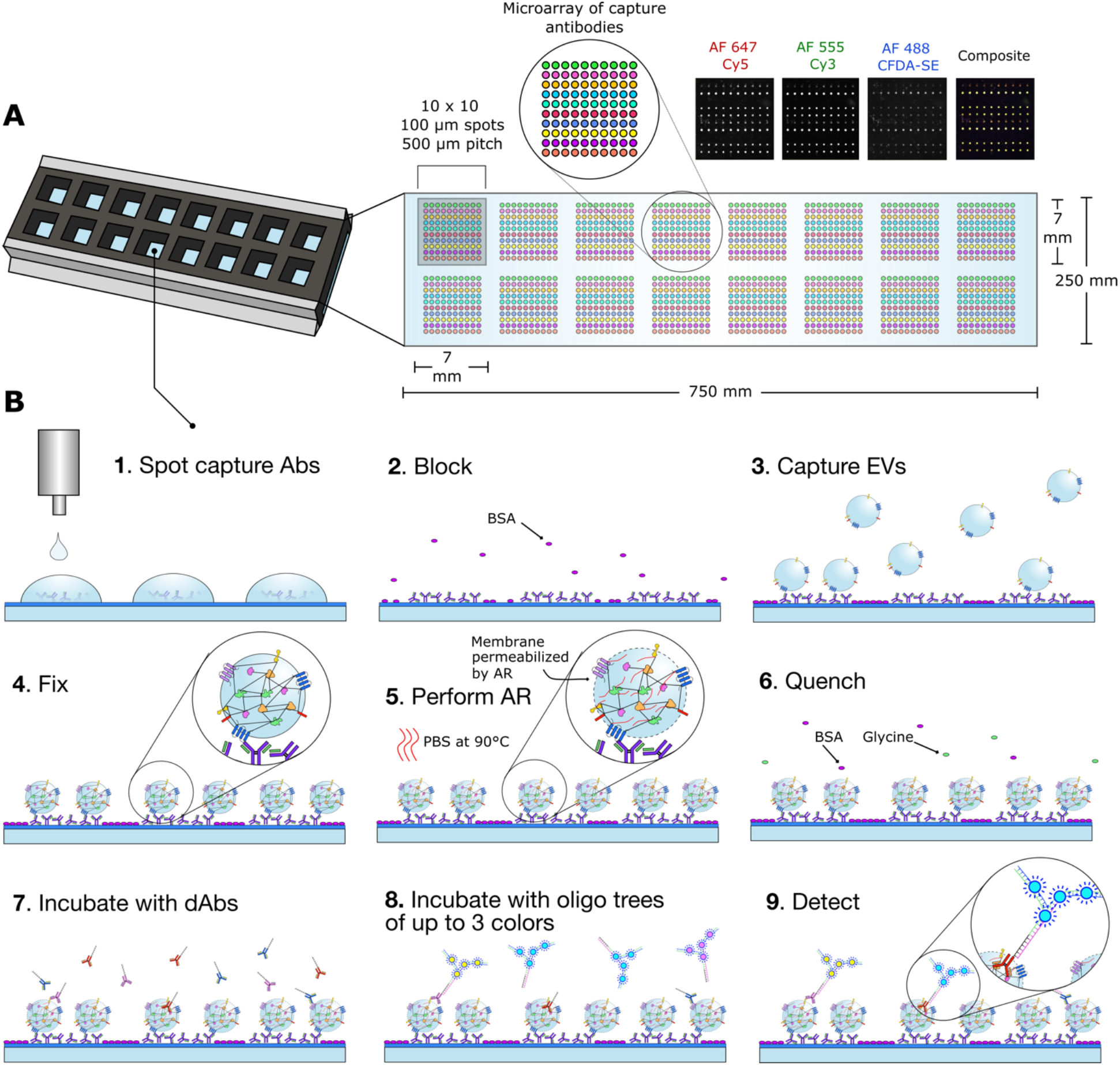
EVPio: combinatorial profiling of extracellular vesicle inner and outer proteins. (**A)** The EVPio assay is performed on a well gasket-covered glass slide inkjet-printed with 16 10×10 subarrays of antibodies, where each row targets a different EV surface protein. (**B)** EVPio workflow: (1) capture antibodies are inkjet-printed on a functionalized slide (PolyAn 2D-Aldehyde), (2) the slide is washed and unoccupied sites are blocked, (3) the sample is incubated on the antibody spots and the EVs are captured based on their surface protein markers, (4) bound EVs are fixed on the slide, (5) the slide is AR-treated and (6) quenched to get rid of remaining reactive groups, (7) the slide is washed and incubated with up to three different oligo barcode-conjugated primary detection antibodies, (8) the slide is washed and incubated with complementary fluorescently-labeled oligonucleotide constructs (up to three colors), and (9) immobilized immunolabeled EVs are detected using a confocal microarray scanner. Microspots, antibodies, oligonucleotides and EVs are not to scale.

### Fixation, Permeabilization and Antigen Retrieval of EVs

The lipid bilayer forms a barrier which must be breached to allow detection antibodies to diffuse inside of EVs and bind inner proteins, while preventing those proteins from diffusing out. Hence, a series of treatments, including fixation with formaldehyde to cross-link diffusible proteins inside the EVs, permeabilization, and AR to make the cross-linked proteins accessible to the detection antibodies, are necessary. Fixation is commonly used for EVs,^39, 40^ can covalently link EVs to the capture antibodies and thus prevent their detachment during subsequent procedures, and does not interfere with the binding of detection antibodies to the outer proteins (data not shown). However, the network of methylene bridges formed following fixation limits immunodetection within the EV lumen by masking binding epitopes and *via* steric hindrance.^33, 41^ In cytochemistry, AR is used to improve detection antibody binding through a mix of hydrolytic breakage of intra- and intermolecular crosslinks and other, buffer-driven effects on protein entangling and renaturation^33^ (Figure 2A). However, permeabilization and AR were found to also affect the detection of outer proteins in EVs, which added a further constraint when optimizing the assay conditions.

**Figure 2.**
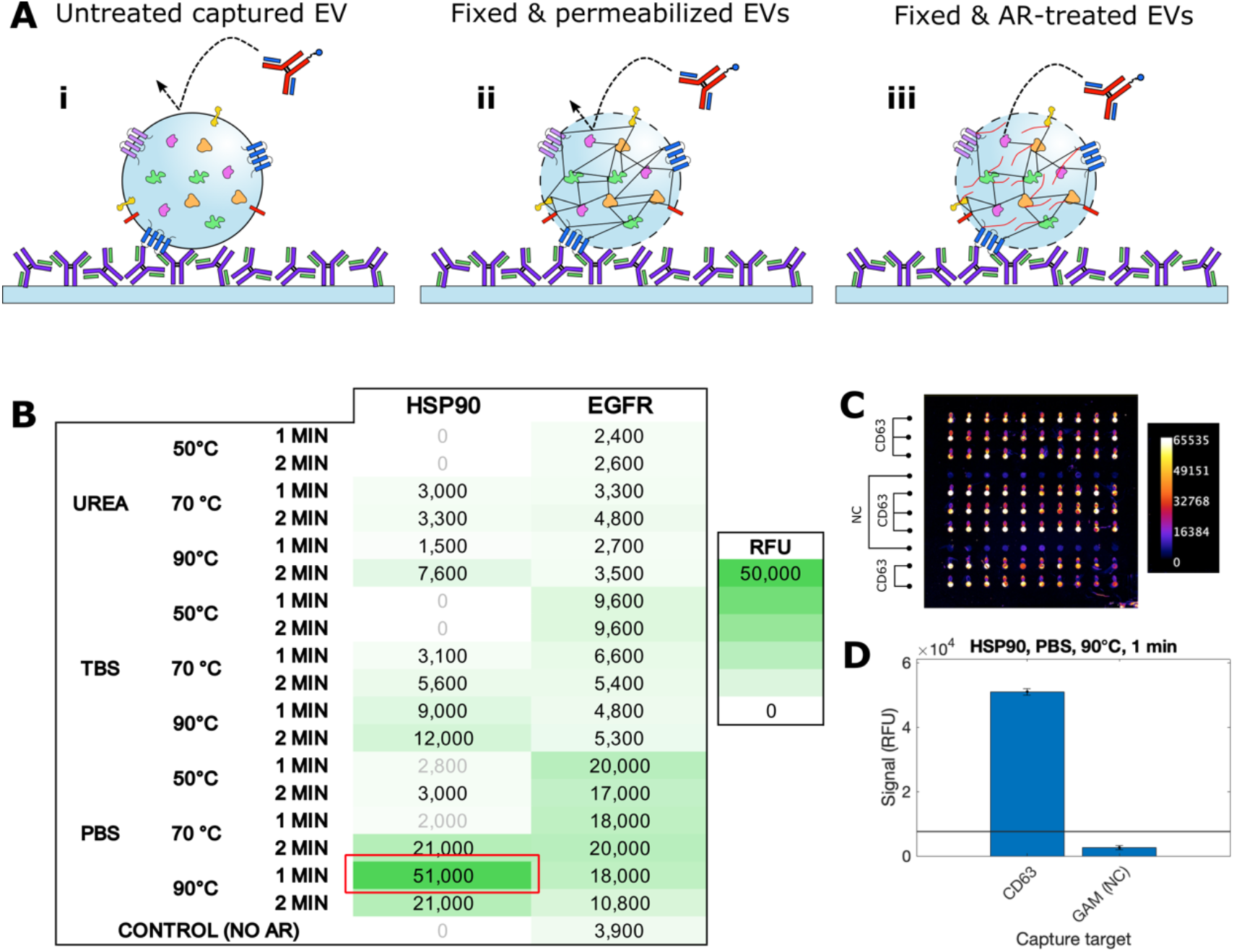
EVPio assay optimization for inner protein detection. (**A)** Illustration of the need for an antigen retrieval (AR) step. (i) The inner proteins of captured EVs are bound by the lipid bilayer membrane and inaccessible to antibodies. (ii) Fixation cross-links inner proteins, while permeabilization allows antibodies to diffuse into EVs, but methylene cross-linking limits accessibility to epitopes. Permeabilization can be used to allow antibodies to breach the membrane while keeping cytosolic proteins in place, but crosslinks can limit epitope accessibility. (iii) Heat-based AR treatment of EVs induces partial hydrolytic breakage of crosslinks, allowing antibody binding of retained and immobilized proteins. (**B)** AR optimization experiments with different buffers/additives, temperatures and incubation times, and fluorescence detection signal for inner protein HSP90 and outer protein EGFR, with superposed heatmap. The red rectangle indicates the optimal condition that was chosen based on high HSP90 and high EGFR signals. Signal values are above the threshold of negative capture control + 2SD unless greyed. A zero value indicates a negative or zero difference between the signal and its associated no EV control. (**C)** Fluorescence micrograph of the optimal AR condition: 1 min at 90°C, in PBS. (**D)** Bar graph of HSP90 detection signal on spots with CD63 capture antibody and negative control spots with goat anti-mouse (GAM) antibody serving as negative capture control (NC). The threshold of negative capture control + 2SD is illustrated as a horizontal line.

AR is a customizable process with many adjustable parameters including buffer, additives, pH, incubation time, temperature, and heating method (microwave, pressure cooker, water bath, *etc*.).^34^ Common additives used in AR include denaturants and chaotropic agents, such as urea.^33^ To adapt AR to EVs, we selected 4 parameters: buffer, temperature, incubation time and presence of a dedicated permeabilization step; and optimized them for simultaneous detection of both HSP90, an inner, cytosolic target, and of EGFR, an outer, membrane receptor tyrosine kinase often overexpressed in cancer.^42^ We used fluorescent signal strength (in RFU) to guide the optimization in order to maximize inner protein detection signal, and manually assessed array images for overall cleanliness. CD63-GFP-expressing EVs from A431 epidermoid carcinoma cells (A431-GFP),^43^ of well-known proteomic composition,^44^ were used as a model and immobilized on the microarray spots using an anti-CD63 capture antibody. A water bath was used for heating, and phosphate buffer saline (PBS, pH 7.0), tris-buffered saline (TBS, pH 9.0), and urea 6 M in PBS (pH 9.5) used as buffers.

The fluorescent signal for HSP90 and EGFR for 18 different AR conditions and a control without AR are shown in Figure 2B. Incubation in PBS at 90°C for 1 min (Figure 2C-D) was observed to give the highest signal for HSP90 and a strong signal for EGFR. The elongated spot shape in Figure 2C is due to comet tails, a phenomenon where minor, directional spreading of the antibody microspots happens upon washing. Fixation followed by permeabilization with various concentrations of Triton X-100,^45^ without AR, allowed to detect inner membrane epitopes of transmembrane proteins, but not free, cytosolic proteins (data not shown). When a separate permeabilization step (10-minute 0.05% Triton X-100) and AR were used, the fluorescence signals were lower than when only AR was used, especially in the case of HSP90. With separate permeabilization, the optimal temperature and time shifted to a cooler, longer incubation (70°C, 2 min in PBS), but only yielded a detection signal of 11,300 relative fluorescence units (RFU) for HSP90, compared to 51,000 RFU for the optimal condition without separate permeabilization (Table S1). These results led us to conclude that AR also permeabilized the EVs and might eliminate the need of a separate permeabilization step. Interestingly, in the absence of dedicated permeabilization with Triton-X, AR with PBS (pH = 7.0) led to more than five times higher HSP90 signals than with TBS (pH = 9.0) or 6 M urea in PBS (pH = 6.0). In tissues, AR is commonly performed in non-neutral pH buffers to generate electrostatic charges that help prevent aggregation upon sample cooling, ensuring that epitopes remain exposed.^33^ However, the effectiveness of this approach is known to be dependent on sample composition and the epitopes probed,^33, 34^ which could account for the differences in optimal protocols. Of note, while the optimal time for AR and heating is 1 min (Figure 2), classic immunohistochemistry protocols typically recommend 10-20 minutes.^33^ Preliminary testing indeed showed that times over 5 min resulted in almost complete loss of signal (data not shown). The significant differences in optimal parameters underscore the differences between EVs and immunohistochemistry imaging of formalin-fixed, paraffin-embedded tissues for which classic AR protocols are optimized. In summary, antibody microarray-bound EVs are more fragile and more easily permeabilized by AR and require much less hydrolytic breakage than tissues, which can be accounted for by the absence of extracellular matrix, cell junctions, and a cytoskeleton, and a different process that omits paraffin embedding and removal.

### Barcoding and Signal Amplification Improve EVPio Assay Performance and Versatility

Multiplexed EV analysis can be accomplished by using multiple detection antibodies with each a different fluorescence spectrum to detect different inner and outer proteins simultaneously. Several options exist to introduce spectral multiplexing. Unlabeled detection antibodies from different species can be recognized by fluorescently labeled secondary antibodies, or by using biotinylated detection antibodies and fluorescent streptavidin together with species-specific secondary antibodies (Figure S1). Whereas for duplex detection these strategies work well, for higher plex, species cross-reactivity becomes a challenge,^46^ as do the constraints and availability of detection antibodies from specific species beyond the most commonly used ones.^47^ Mixing biotin-streptavidin and species-based detection introduces variation through the differences in the way targets are detected, requiring additional adjustments to the data before it can be interpretated. The use of oligonucleotide barcodes to label antibodies in multiplexed settings is often used to address the aforementioned issues^48, 49^ and was adopted to introduce 3-plex detection on EVPio microarrays.

To achieve the signal amplification needed to detect inner proteins, we introduced branched DNA signal amplification and evaluated oligonucleotides with linear (Figure 3A), 2-branch (Figure 3B) and 4-branch (Figure 3C) tree designs with increasing numbers of fluorophores. Linear oligo detection involves antibodies each labeled with a linear, 15-mer barcode, an imager strand containing a barcode-complementary section and an additional barcode for fluorescent probe hybridization, and a complementary probe strand conjugated to a fluorophore (Figure S10). Barcodes were designed to be orthogonal to avoid cross-talk. The 2-branch and 4-branch trees contain additional, intermediate strands that form branches, which are pre-assembled using unique intermediate barcodes that were also designed to minimize cross-reactivity between strands. The 2-branch and 4-branch trees have the capacity to hybridize 4× and 8× more fluorescent probe strands and have the potential to improve assay signals by those factors. However, accessibility of inner proteins to larger trees, different binding dynamics, and steric hindrance may constitute trade-offs of larger trees, and result in a smaller number of binding events and thus lower signal. The optimal design was thus evaluated experimentally.

**Figure 3.**
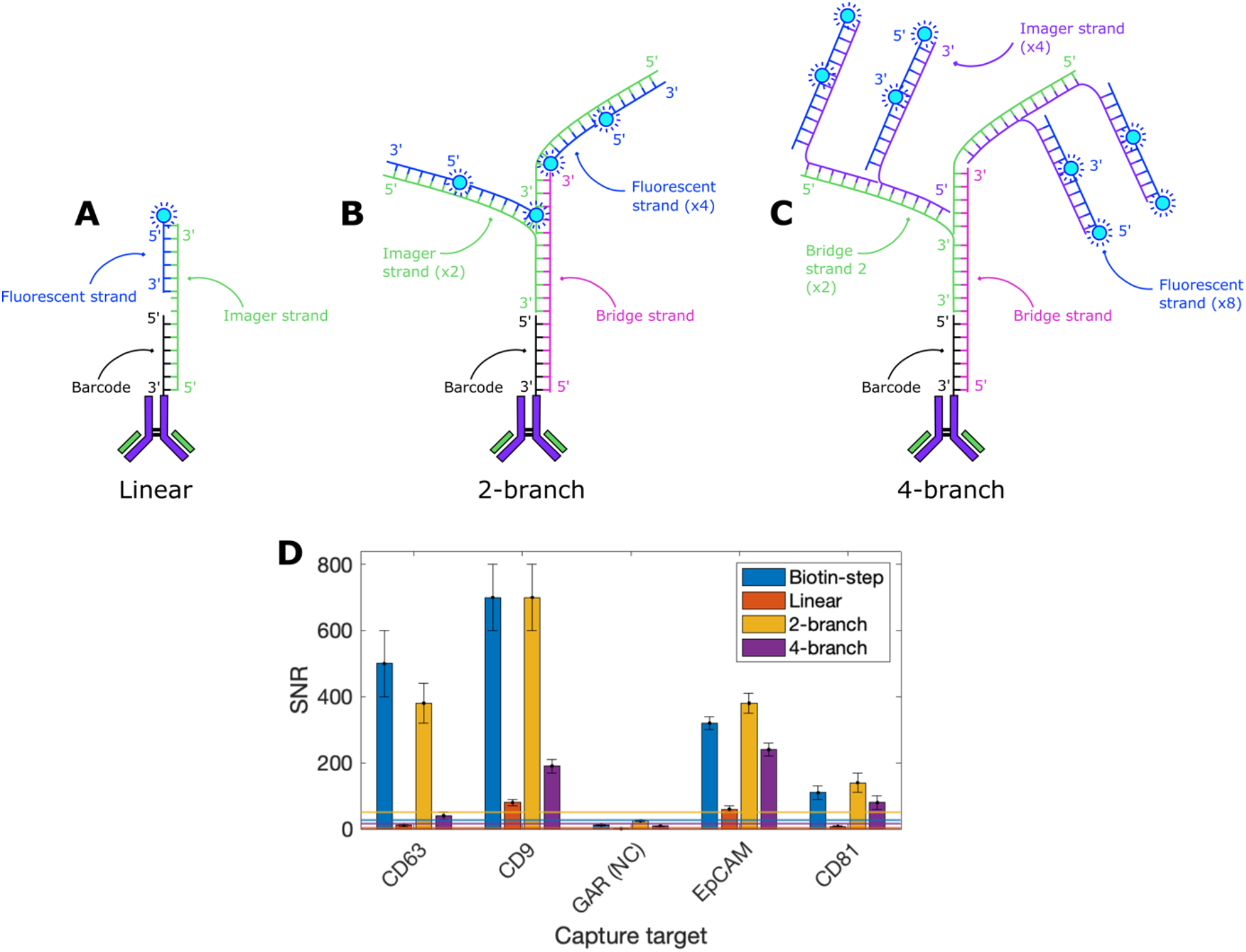
Branched DNA amplification improves EVPio assay performance. Schematics of oligonucleotide constructs tested for signal amplification: (**A**) linear strand with one fluorophore; (**B**) 2-branch tree with 4 fluorophores; (**C**) and 4-branch tree with 8 fluorophores. Length ratios of oligos are not to scale. (**D)** SNR of biotin-streptavidin, linear, 2-branch and 4-branch detection schemes for CD9 detection of SW403 EVs. Barcoded antibodies were first incubated and bound to the proteins expressed in immobilized SW403 EVs, followed by incubation and hybridization of pre-assembled oligo trees bearing Alexa Fluor 647 fluorophores. The 2-branch amplification design gave signals closest to the benchmark biotin-streptavidin detection. Signal thresholds for each detection scheme, calculated as the negative capture control (GAR) signal + 2SD, are indicated by horizontal lines color-matched to the corresponding bars. Error bars are SE.

We used signal-to-noise ratios (SNRs) to guide the optimization of the EVPio assay protocol for oligonucleotide-based labeling and signal amplification, including washing protocol, quenching solutions, incubation and hybridization buffers, and combinations of specific additives such as sheared salmon sperm DNA (SSS DNA) and dextran sulfate (DS) for linear, 2-branch and 4-branch amplification (Figures S2 and S3). Following optimization, again using CD9 detection, this time following capture with antibodies against EV-associated tetraspanins CD9, CD63 and CD81 and adhesion molecule EpCAM, we benchmarked linear, 2-branch and 4-branch amplification (without AR) against biotin-streptavidin detection in EVs from SW403 cells (Figure 3D). The SNRs obtained with biotin-streptavidin detection and the 2-branch construct were statistically comparable for all EV subpopulations (p > 0.05), while linear detection and 4-branch amplification led to significantly different SNRs (p < 0.05), except in the case of the CD81+ subpopulation detected with the 4-branch tree (p = 0.29) (Figure 3D). These results, which echo those obtained in the aforementioned assay optimization experiments, may reflect the increased steric hindrance between 2-branch and 4-branch trees. We adopted the 2-branch design for subsequent experiments as a trade-off between having the maximal signal and minimizing possible interference due to steric hindrance, a factor that becomes even more important with the introduction of multiplexing.

### Multiplexed Analysis of Inner and Outer EV Proteins

The adoption of branched oligo trees for multiplexed EVPio analysis was first explored and optimized with duplex two-color (Alexa Fluor 488 and Cy5) detection of HSP70 as the inner protein and CD9 as the outer protein, using four different capture antibodies (Figure S4). Using again CD63, CD9, EpCAM and CD81 as capture antibodies, we found that 2-branch trees were optimal. We further verified whether multiplexing might skew the signals by comparing them to those obtained through a singleplexed analysis carried out in parallel, and found them to be concordant for EVs from SW403 cells, while for HT29 EVs, multiplexing resulted in signal loss for CD63 detection, regardless of amplification, and gain for CD9 and HSP70 detection. The same analysis was conducted for triplex detection with the simultaneous detection of HSP70, CD9 and CD63 with Alexa Fluor 647, Cy3 and Alexa Fluor 488, respectively, both for HT29 (Figure 4), and SW403 (Figure S5). In this series of experiments, triplexing (compared to singleplex) of protein detection led to signal increases of 2.9 on average (median of 1.4) for EVs from HT29, and 3.9 (median 2.9) for EVs from SW403 cells.

**Figure 4.**
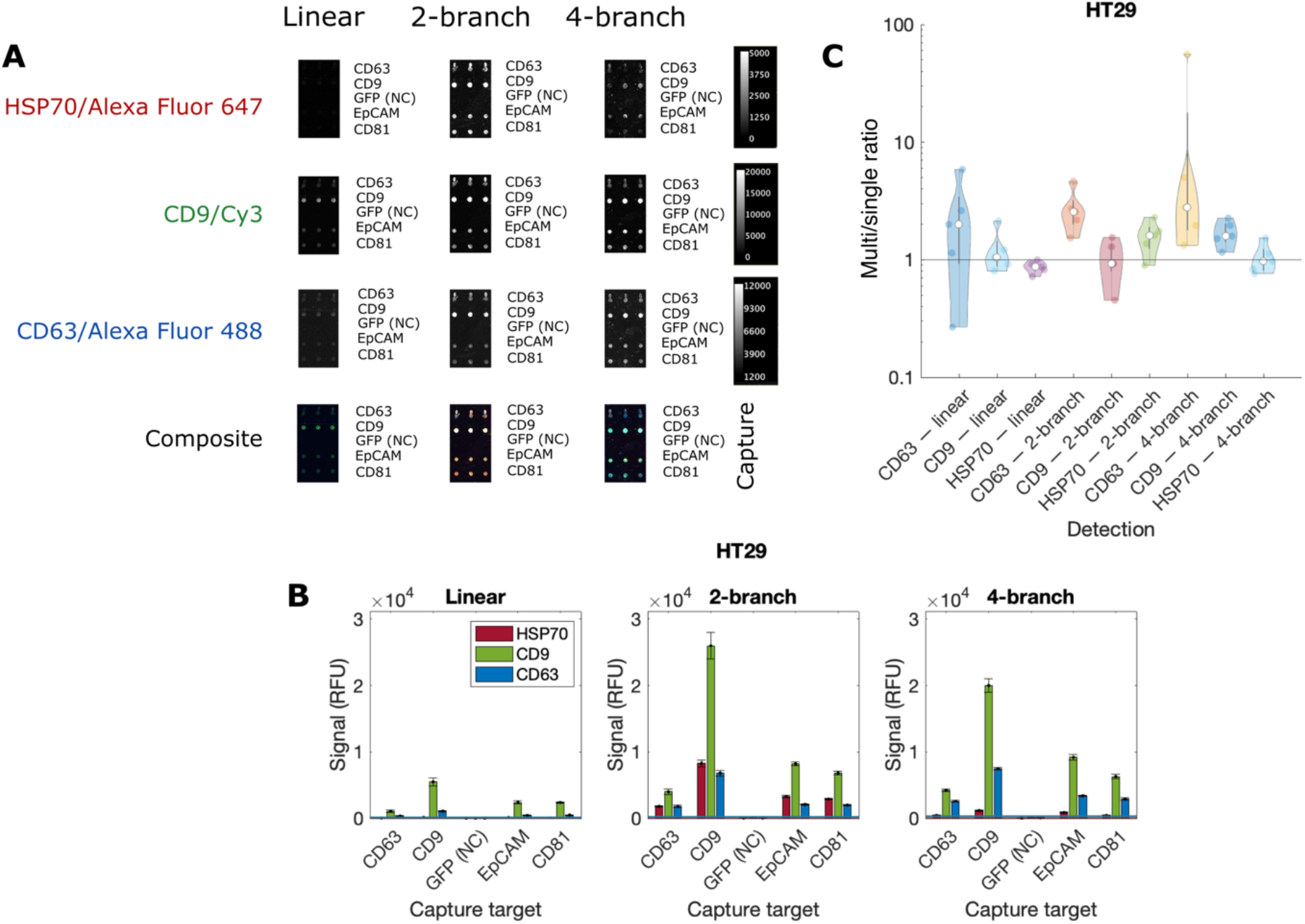
Combinatorial EVPio analysis of inner, cytosolic proteins and outer, membrane proteins and validation of multiplexing through multiplexed *vs* singleplexed analysis. **(A)** Fluorescence micrographs of inner protein HSP70 and outer proteins CD9 and CD63 detected on EVPio microarrays with CD63, CD9, CD81 and EpCAM capture spots, and negative controls (GFP) in HT29 EVs. Complete array images can be found in Figure S6. **(B)** Bar graphs of linear, 2-branch and 4-branch trees for signal amplification. 2-branch amplification gave the highest signals. Errors bars are SE. **(C)** Violin log plots of the averaged ratios for all 4 capture antibodies (CD63, CD9, EpCAM and CD81) as well as a non-specific control (GFP) of multiplexed over singleplexed signals for inner (HSP70) and outer (CD9, CD63) detection antibodies and different amplification trees for HT29 EVs. Averaged multiplexed signals are comparable, and up to ∼ 2.5× higher than singleplexed signals. Violin plots are used to show the distribution of the data, and overlay a boxplot (median, interquartile range) with a kernel density plot (probabilities).

In HT29 and SW403 EVs, inner and outer proteins were simultaneously detected in all four subpopulations with signal amplification (Figures 4A-B and S5A-B). HSP70 could be detected consistently above threshold with 2- and 4-branch amplification constructs, with 3.5 – 13 x higher signals for 2-branch amplification, reaching 10,200 RFU in the SW403 CD9+ EV subpopulation. Inner targets, which are cross-linked to surrounding proteins within fixed and AR-treated EVs, may be more accessible to 2-branch constructs. These results show that it is possible to simultaneously detect outer and inner proteins such as HSP70 with EVPio analysis and 2-branch amplification.

In summary, singleplexed and multiplexed values are broadly in agreement, with multiplexing occasionally producing higher signal and only very rarely signal loss (Figures 4C and S5C). In particular, 2-branch-based signal amplification gave signals that agree and even improve upon singleplexed values, especially for HSP70 detection. These findings strengthened our decision to use the 2-branch design for further EVPio assays.

### EVPio Analysis Recapitulates Key Proteomic Characteristics of EVs from CRC Cell Lines

To showcase the potential of EVPio analysis, we phenotyped inner and outer proteins in EVs from CRC cell lines HT29 and SW403. EVs were captured on the array by targeting the three ‘canonical’ EV-associated tetraspanins (CD9, CD63, CD81)^50^ or EpCAM, an adhesion molecule overexpressed in epithelial cancers.^51^ A panel of 12 protein targets were selected for their expression in EVs and their biological function. The outer proteins were all membrane proteins, and include the three tetraspanins used for capture (CD9, CD63, CD81) as well as CD82, another tetraspanin and known tumor metastasis suppressor;^52^ and the integrin (ITG) subunits (α2, α6, β1, β4)^8^ that have been associated with EV-mediated metastatic organotropism.^8^ Among the inner proteins was the tight junction protein claudin-2,^53^ a prognosis biomarker of replacement-type liver metastasis in EVs from CRC patients,^54^ detected through its cytoplasmic domain, and cytosolic proteins, which were selected based on known and widespread association with EVs.^55^ They included the heat-shock proteins (HSP90, HSP70),^56^ and ESCRT-associated protein Alix, which is involved in EV biogenesis and cargo sorting.^2, 57^ Targets were divided in 4 sets and used for triplexed detection (Figures 5A-B and S7). The inner and outer protein profiles are shown in Figure 5C-D. Figure 5C shows the absolute RFU values of fluorescence expression for HT29 EVs, while Figure 5D shows the relative SW403/HT29 expression for comparison. This analysis protocol using one set as a reference allows to highlight differences, avoids the need for a binding curve or calibration (which unlike for ELISA are not used in EV analysis to date), and could help distinguish between healthy and diseased samples, for example. The condition however is that all samples be run as one experiment under the same conditions to avoid introducing experimental errors. To evaluate the reproducibility of our approach, a replicate experiment was performed, and the two measurements compared (Figure S8). The results from the two repeat experiments were highly correlated.

**Figure 5.**
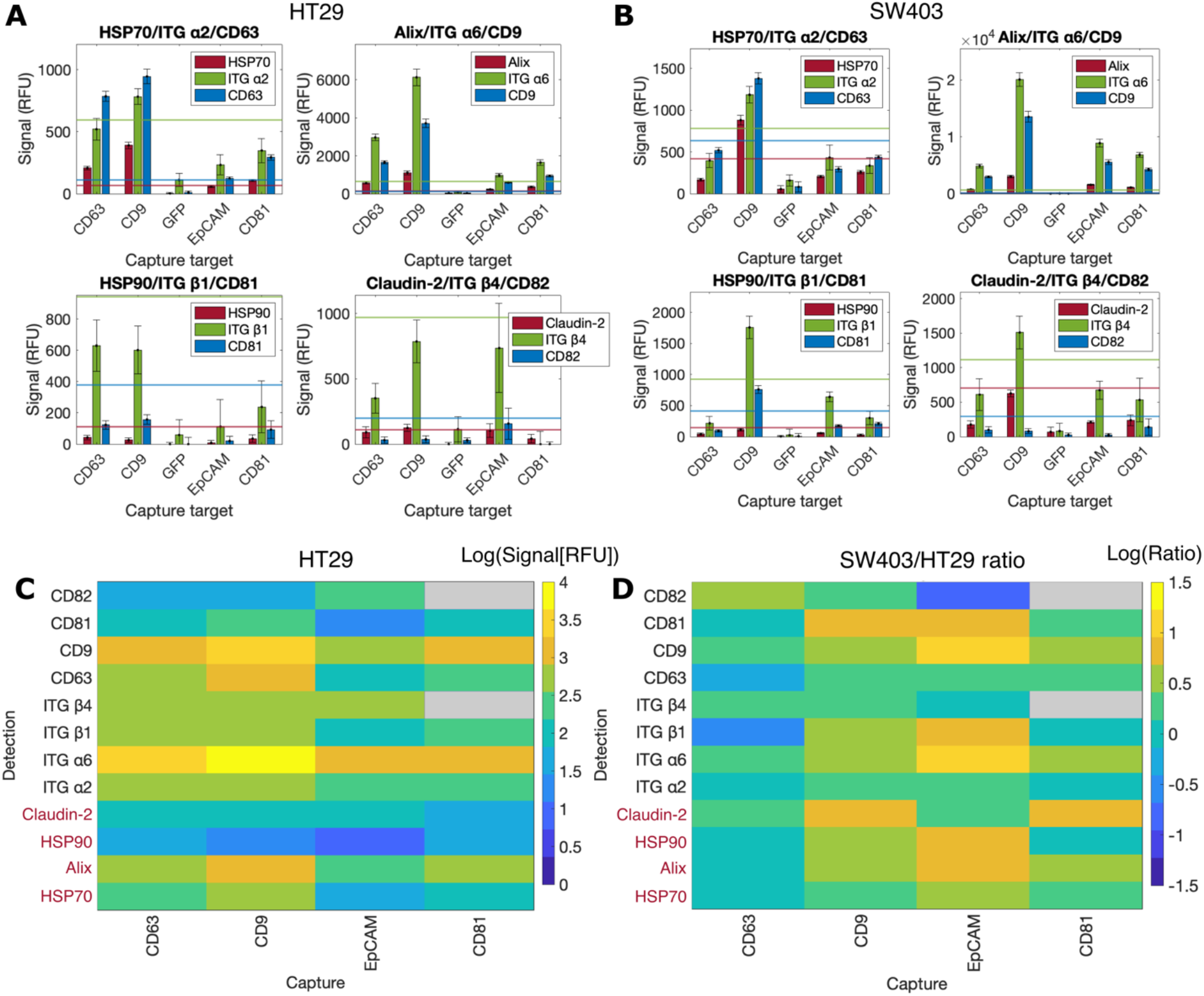
Multiplexed EVPio analysis of HT29 and SW403 EVs immobilized with 4 capture antibodies and a negative capture control (GFP), with 4 separate 3-plex assays to detect 12 inner and outer proteins. Assay signals in (**A**) HT29 and (**B**) SW403 EVs. Bar graphs are color-coded to reflect the fluorophore/channel used for each target (red for inner proteins, green and blue for outer proteins), with each separate graph representing a 3-plex assay. Detection thresholds for each target are calculated from anti-GFP negative control + 2SD and indicated by horizontal lines color-matched to the corresponding bars. Error bars are SE. Multiplexed phenotyping results for **(C)** HT29 and **(D)** SW403 EVs. The values for SW403 EVs are expressed as the ratio of the SW403 to the HT29 signal, highlighting differences between EVs from the two cell lines. Inner proteins are colored in red for emphasis. In HT29 EVs, the CD9^+^ EV subpopulations presented the highest overall signals, with ITG α6, CD63, ITG β1 and ITG β4 as notable detected markers. HSP70 and Alix were the strongest inner targets. SW403 EVs presented notably higher ITG α6, CD9 and CD81 signals in the EpCAM^+^ EV subpopulation. Greyed values represent absence of signal.

Among the 4 capture antibodies, the CD9 subpopulation led to strong overall detection signals, consistent with a high number of vesicles captured. CD9 detection signals were strong as well and exceeded the threshold of the detection signal for the negative capture control (anti-GFP) plus 2 SDs in all cases, with the highest signal found in the CD9^+^ subpopulation of SW403 EVs. The fluorescence for other tetraspanins was lower, and in some cases did not exceed the threshold. As can be seen in several panels, the background noise measured on the anti-GFP antibody spots is significant and explains why signals that are clearly visible are below the threshold (also see discussion below). ITG subunit α6 was the protein marker with the highest detection signal across all captured EVs, reaching 20,000 RFU and 9,000 RFU in CD9^+^ and EpCAM^+^ SW403 EVs, consistent with the over-expression of ITG α6β4 in colorectal cancer.^58^ ITG β1, whose high expression in CRC is associated with poor prognosis,^59^ was notably detected in CD9^+^ SW403 EVs, but was also found at values close to threshold in EpCAM^+^ SW403 EV as well as CD63^+^ and CD9^+^ HT29 EVs. Inner proteins, including both untethered cytosolic proteins as well as the inner leaflet domain of trans-membrane proteins, could also be detected. Alix gave consistently above threshold signals in all EV subpopulations under study, reaching a maximum of 1,100 and 3,000 RFU in HT29 and SW403 EVs, respectively, in accordance with its involvement in the ESCRT pathway and EV biogenesis.^2, 57^ While HSP90 was not detected above threshold, notable signal was obtained for the CD9^+^ SW403 EV subpopulation. Claudin-2 was primarily detected in CD9^+^ EV subpopulations of both cell types, with additional notable expression in CD63^+^ and EpCAM^+^ HT29 EVs, while HSP70 gave signal in all subpopulations, with CD9^+^ EVs of both types and CD63^+^ HT29 EVs yielding above threshold values.

Our data shows that despite the genetic differences of their originating cells, HT29 and SW403 EVs present similar overall profiles regarding the 12 proteins included in our proof-of-concept panel, albeit with some key differences (Figure 5D). In SW403 EVs, the EpCAM^+^ subpopulation presented approximately 10× higher signals for CD81, CD9 and ITG α6, while inner proteins HSP90 and Alix had ∼6× stronger signals compared to HT29 EVs. The CD9^+^ and CD81^+^ subpopulations also had higher signals (∼3-5×) for several detection targets, including CD81, CD9, ITG α6, and claudin-2. Notable targets with lower signals included CD82 in the EpCAM^+^ subpopulation and ITG β1 in the CD63^+^ subpopulation. The CD63^+^ subpopulation had the largest number of lower signal targets, with nearly half presenting lower signals than in HT29 EVs.

A caveat of assessing inner and outer targets simultaneously post-AR is loss of signal for outer targets, which can be expressed as the ratio between the signals obtained with and without an AR step (Figure S9). While AR usually enhance inner protein signals (when inner protein detection can be achieved pre-AR), with an average AR/no AR ratio of 4.1, outer proteins signals are usually reduced, with an average ratio of 0.25. This signal loss can potentially be avoided for smaller target subsets through optimization of the process to the specific antibodies used,^34^ but performing high-throughput and multiplexed analysis for a panel of proteins requires a single, one-size-fits-all AR condition.

In summary, the EVPio platform can be used to profile the protein content of EVs originating from different cell types in a multiplexed and high-throughput manner.

## Discussion

Here, we introduce the EVPio analysis platform that includes an antibody microarray, EV-specific AR treatment, antibody barcoding, and branched DNA signal amplification at the labeling step for the multiplexed analysis of inner and outer protein co-expression patterns in EVs *in situ*. On a single slide of 16 arrays incubated each with 50 μL of sample, we quantified 12 proteins in EVs from two cell lines captured using 4 different capture antibodies, along with controls. Following extensive optimization of fixation and AR, a simple protocol was obtained which only adds 1 min incubation in PBS at 90 °C after fixation and simultaneously permeabilizes EVs and restores antigens, thus enabling simultaneous probing of inner and outer proteins. Of note, the signal intensity for the outer proteins was lower after AR. Hence, depending on the application, and notably if there is a desire to compare the results to samples that did not undergo AR (nor permeabilization), it may be advantageous to incubate the sample on two replicate sets of microarrays, and use one set for multiplexed outer protein detection, and the other, with AR, for multiplexed inner protein detection. The use of oligo-conjugated antibodies and 2-branch DNA trees (see also further below) eliminates many constraints on detection antibody usage while fluorophores can be varied from experiment to experiment by changing the fluorescent probe oligo.

The steps of the EVPio assay were optimized sequentially as the workflow was constructed, using parameters and ranges narrowed down based on the literature and preliminary experiments. The initial optimization of the AR process was performed using fluorescently labeled secondary antibodies and primary antibodies against model proteins HSP90 and EGFR, and only a subset of possible incubation times and temperatures were tested to limit the number of experiments (Figure 2B and Table S1). The resulting optimal parameters were retained after branched DNA amplification was added to address low inner protein signals and enable multiplexing without species constraints using combinations of oligo-labeled detection antibodies. It is thus possible that assay conditions could be further optimized, for example through testing using the proteins to be analyzed and branched DNA amplification. Design of experiments (*e*.*g*. Taguchi method) can help explore a large range of factors including buffer compositions, surfactants, temperatures, incubations times and signal amplification, and both identify the optimal factor values and quantify their respective contribution to the chosen optimization parameter, *e*.*g*. assay SNR.^60, 61^ Whereas here optimization was conducted based on the expression of one outer protein, the inclusion of multiple inner and outer proteins during the optimization of the AR protocol could help identify broadly applicable, optimal trade-offs between inner and outer protein detection.

Reproducibility and quality control are central to reproducible experiments and science, and both reagent-driven cross-reactivity and spatial bias will need to be considered. Reagent-driven cross-reactivity can arise when mixing multiple detection antibodies^62^ and lead to false positive signals, such as the ones observed here with our negative anti-GFP control in some experiments. We previously identified spatial bias as a likely underestimated source of variability in antibody microarrays,^63^ and it is of particular concern when multiple samples are measured at different locations on a slide. We performed a visual, qualitative check of each functionalized slide prior to the assay by scanning pristine slides with strong signal gain in the 532 nm channel. The slight autofluorescence of the aldehyde coating produced a visible signal and slides with immediately visible irregularities were discarded. The possible impact of spatial bias on EV microarrays assay reproducibility and variability was not studied in depth, however.^63^

The addition of oligonucleotide barcoding circumvents common multiplexing issues such as species cross-reactivity and limited availability of species or directly labeled antibodies. The best suited primary antibody can be used for every target, independently of its species, clonality, or availability as a conjugate. Barcoding also offers flexibility for future integration with higher-multiplexing schemes, such as next-generation sequencing^64, 65^ or DNA exchange imaging,^48^ and alternative signal amplification methods, like polymerase chain reaction (PCR) or rolling circle amplification (RCA).

Branched DNA signal amplification was explored to improve assay performance while minimizing assay time and cost. The oligo trees, which can be assembled in parallel to the main assay, can directly be incubated on the arrays, circumventing the need for enzymes, heating and temperature control, and are easily customized with any fluorophore label. Furthermore, the size of the trees can be selected prior to the assay based on the expected abundance of the target protein and the associated anticipated level of steric hindrance to improve signal strength (Figure 3). For low target concentrations, there would be enough space for antibodies to bind with large trees, while at high concentrations, the binding of one antibody and a large tree could limit binding to adjacent sites by other antibodies conjugated to large trees, thus leading to better signals for smaller trees. In addition, as up to three proteins are detected simultaneously, steric hindrance effects could become more acute with higher multiplexing, which solidified our choice of the intermediate 2-branch tree.

As a proof of concept, we showed the potential of EVPio analysis for phenotyping and comparing EV samples from two CRC cell lines. Using a panel of 4 different EV capture antibodies combined with 12 different protein detection antibodies including tetraspanins, integrins, a disease-related claudin and EV-associated cytosolic targets, we showed how EVPio can be used to characterize and compare EVs. The protein co-expression patterns that were identified were consistent with the CRC origin of the EVs, revealing both similarities and some differences between the two originating CRC cell lines. While the SW403/HT29 expression ratio of the majority of pairwise target combinations assessed were within a factor 3×, some co-expression patterns stood out as up to 10× higher (CD81, CD9 and ITG α6 in EpCAM^+^ EVs) or lower (CD82 in EpCAM^+^ EVs, ITG β1 in CD63^+^ EVs) in SW403 EVs. Certain targets did not meet the threshold of two standard deviations above negative capture control (anti-GFP) spots background. However, non-specific binding between the capture and detection antibodies, and residual fixative-induced autofluorescence (despite quenching) on that spot meant that cutoff threshold values increased. The 20 replicates used here helped map the variability, and establish the high thresholds found in the Cy3 fluorescence channel, which in our setup is the emission filter with the largest bandwidth, rendering it particularly susceptible to the aforementioned sources of background fluorescence. Strategies to improve the assay accuracy and lower thresholds include replacing the Cy3 by another fluorophore with emission that peaks further away from the autofluorescence peak (such as near-infrared fluorophores) and/or switching to an emission filter with a smaller bandwidth to minimize the contribution of autofluorescence, at the possible cost of signal sensitivity.

A trade-off to using different fluorophores are the different signal intensities in each channel, which make quantitative analysis of protein expression challenging. It might be possible to develop a normalization protocol to account for differences between the fluorophores (brightness) and imaging setups (lasers, filters, *etc*.) used for each fluorescence channel, but variable backgrounds, and low signals, may compromise quantitative analysis.

Given the recent popularity of EVs and their potential for understanding health and disease, many existing technologies have been harnessed and new platforms created to study the EV proteome, including ensemble average techniques such as mass spectrometry, custom microfluidic devices, affinity binder arrays, and commercial systems like ExoView. EVPio stands out with its blend of high customizability (any antibody can be used, from any species), combination of inner and outer protein detection with high antigen retention, effective signal amplification, and compatibility with common lab equipment, including either a microarray scanner or fluorescence microscope.

In summary, we described an antibody microarray-based workflow for the simultaneous detection of transmembrane and cytosolic proteins in EVs, which can be applied to phenotype EVs and derive co-expression patterns in a multiplexed and high-throughput manner. We detailed the optimization of critical aspects—inner protein detection, multiplexing and signal amplification— and showcased how EVPio analysis can be used to uncover disease-relevant protein co-expression patterns in EVs from cancer cell lines. The loss of signal of outer protein expression following AR should be considered when designing an experiment. Depending on the application, inner outer protein may be measured simultaneously on the same slide, or on separate slides with only the ones for inner protein analysis subjected to AR. EVPio analysis opens the way to extensive combinatorial protein profiles with significance for EV characterization and biomarker research in health and disease.

## Materials & Methods

### Cell Culture

CD63-GFP-transfected A431 cells (ATCC® CRL-1555TM, provided and transfected by Dr. Janusz Rak, McGill University, Montreal, Canada), HT29 cells (ATCC® HTB-38) and SW403 cells (ATCC® CCL-230) were cultured in Dulbecco’s Modified Eagle Medium containing 4.5 g/L D-glucose, L-glutamine and 110 mg/L sodium pyruvate (DMEM, Gibco) and Rosewell Park Memorial Institute 1640 (RPMI 1640, Gibco) media containing L-glutamine, respectively, and supplemented with 10% fetal bovine serum (FBS, Gibco) and 1% penicillin-streptomycin (PS, Thermo Fisher Scientific). Cells were incubated at 37°C and 5% CO_2_ with constant humidity.

### EV Preparation and Characterization

Cells were passaged into T75 flasks (Corning) for expansion. 24h after seeding, culture media was replaced with new media supplemented with 5% exosome-depleted FBS (Gibco) and 1% PS. Once the cells reached 80% confluency, the cell media was harvested into 50 mL tubes, spun at 400 g for 15 min to pellet cell debris, syringe-filtered (pore size 0.22 μm, 33 mm diameter, MilliporeSigma) and concentrated down to 500 μL per 30 mL through iterative 25-min spins at 4000 rpm in Amicon Ultra-15 centrifugal filters (100 kDa cutoff; Millipore Sigma). Size-exclusion chromatography (SEC) was performed using qEVoriginal columns (70 nm; Izon Science). Each column was loaded with 500 μL of concentrated sample, then flushed with filtered PBS (0.22 μm filter, MilliporeSigma) while the eluate was collected in ten 500 μL fractions.

A NanoDropTM 1000 spectrophotometer (Thermo Fisher Scientific) in A280 mode was used to estimate the protein content (mg/mL) of the fractions. Fractions 8 and 9 of each set were pooled and the concentration (particle/mL) of the resulting sample assessed by Tunable Resistive Pulse Sensing (TRPS) (qNano, Izon Science).

### General EVPio Assay Workflow

Antibody microarrays were patterned on PolyAn 2D-Aldehyde slides (PolyAn) using the sciFLEXARRAYER SX inkjet bioprinter (Scienion). Slide were imaged in the 532 nm channel at 100% gain prior to patterning to qualitatively assess the uniformity of the aldehyde functionalization. Antibodies were diluted at 100 μg/mL in a solution of 15% 2,3-butanediol (Sigma-Aldrich) and 1 M betaine (Sigma-Aldrich) in PBS and spotted in 16 (2 × 8) subarrays of 100 (10 × 10) 100-μm spots with a 500 μM pitch. Patterned slides were incubated overnight at 70% humidity, washed for 15 min in 0.1% Tween-20 (T20, ThermoFisher) in PBS, blocked for 3h in a solution of 3% BSA (Jackson ImmunoResearch) and 0.1% T20 in PBS, dried, and inserted into 16-well gaskets (Grace Bio-Labs).

A portion of each EV sample was dyed for validation purposes through incubation with CFDA-SE (V12883, Invitrogen) at 1/250 concentration for 2h at 37°C. Dyed and undyed samples were then supplemented with 1% BSA and incubated with mild agitation (450 rpm) over the arrays for 2h at room temperature followed by overnight at 4°C. Samples were washed with washing buffer (0.03% T20 in PBS) at 450 rpm, then fixed in 4% formaldehyde (diluted in PBS from 16% methanol-free formaldehyde (w/v) solution, ThermoFisher). For the antigen retrieval step, gaskets were removed and arrays were incubated for 1 min in 90°C PBS. Following a MilliQ water wash, gaskets were reapplied and the arrays were quenched for 2h at room temperature (450 rpm) in a solution of 3% BSA, 0.3 M glycine and 0.03% T20 in PBS. Detection antibodies at a concentration of 1 μg/mL (outer targets) to 2-5 μg/mL (inner targets) were incubated for 2h at room temperature (450 rpm) in incubation buffer (3% BSA, 0.03% T20 in PBS).

Gasket wells were washed 3 times in washing buffer, then labeling was performed either with fluorescent streptavidin/fluorescent species-matched secondary antibodies (both 1:1000 in incubation buffer) or fluorescent oligonucleotide probes/constructs (10-50 nM in PBS with 0.025% T20, 300 mM NaCl, 0.5 mg/mL sheared salmon sperm DNA [SSS DNA, ThermoFisher] and 0.5% dextran sulfate [Sigma]) for 30 min at room temperature, 450 rpm. When using oligonucleotides for detection and labeling, the strands composing each amplification construct (see Figure 3A-C) were mixed and annealed using a Biometra T-Gradient thermocycler prior to labeling, and blocking, quenching and incubation buffers were supplemented with 0.5 mg/mL SSS DNA. The slides were then washed for 3 min in washing buffer, dried, and imaged using a confocal microarray scanner (InnoScan 1100 AL, Innopsys). For antibody or streptavidin-based labeling, slides were washed for 15 min and rinsed with MilliQ water prior to drying and imaging.

### Optimization of Inner Protein Detection

A431-GFP EV samples were captured on microarrays of anti-CD63 (353014, Biolegend) and goat anti-mouse (A16080, Invitrogen) antibodies and fixed as described in the previous section. At the antigen retrieval step, 19 combinations of buffer (6 M urea [Sigma] in TBS [Fisher], pH 9.5; TBS; PBS) temperature (50°C, 70°C, 90°C), incubation time (1 min, 2 min) and permeabilization status (with or without incubation in 0.05% Triton-X [Sigma] in PBS for 10 min) were tested (Figure 2B). Claudin-1 (5190000, Life Technologies), HSP90β (PA3-012, Invitrogen) and biotinylated EGFR antibodies (BAF231, R&D Systems) were used at the detection step at concentrations of 5 μg/mL, 5 μg/mL and 1 μg/mL, respectively, followed by labeling with Alexa Fluor 647-conjugated goat anti-rabbit antibodies (A-21245, Invitrogen) and streptavidin (S21374, Invitrogen).

### Barcoding and Amplification Tree Design

Figure 3A-C list the oligonucleotide strands used for amplification as well as the names of their constituting sequences. Sequences are detailed in Figure S10. All oligonucleotides were supplied from IDT.

For the linear probe design, the unique 21-mer fluorescent barcode (FBC) and a unique toehold linker were designed sequentially by using EGNAS^66^ to generate libraries of sequences, with FBC used as neighbor for the linker library. The resulting sequences were checked for secondary structures with QuickFold.^67^ Orthogonal 15-bp nucleotide barcode (BC) sequences were obtained by using EGNAS and PrimerPooler^68^ for sequential sequence generation and cross-reactivity verification, respectively, using FBC and the linker as neighbors, barcodes from previous runs as included sequences, and ΔG = -7 as a cross-reactivity threshold. NUPACK^69^ was used to assess barcode interaction and narrow down the sequences to a smaller number of orthogonal barcode sequences.

For the intermediate barcode (IBC, 2-branch tree), EGNAS^66^ was used to generate a set of 21-mer sequences with 40-60% GC content, low hairpin tolerance, the complements of the orthogonal barcodes and the fluorophore barcode as neighbors, and quadruple of any nucleic acid as forbidden sequences. The resulting sequences were fed to PrimerPooler^68^ to screen for cross-reactivity (threshold of ΔG = -6) against the orthogonal barcodes, the fluorophore barcodes, and their complements. The remaining sequences were checked for folding patterns as part of the complete bridge and imager strands with QuickFold,^67^ and a sequence with melting temperatures not exceeding 50°C was chosen. An analogous pipeline was used for the design of the 4-branch tree (IBC1 and IBC2). For IBC1, EGNAS was used as previously but without neighbors, and ΔG was set to -5.35 in PrimerPooler. For IBC2, IBC1’ and FBC’ were set as neighbors, and cross-reactivity was checked against all BCs and BCs’; IBC1 and IBC1’; and FBC and FBC’. QuickFold was used to check for folding within the bridge 2 and imaging strands.

### Antibody Conjugation with Oligonucleotide Barcodes

Antibodies to be conjugated were supplemented with a 30× molar excess of sSMCC (ThermoFisher), diluted down to a concentration of 0.5-2 mg/mL in PBS, and left to for 2h at room temperature. Oligonucleotides BCs with a /3ThioMC3-D1-3’ modifier, in a 9× molar excess to their corresponding antibodies, were supplemented with 50-200 mM DTT (ThermoFisher) and diluted to 20-100 μM in PBS. The mix was incubated for 2h at 37°C under mild agitation (150 rpm). Purification was performed with 7kDa cutoff Zeba Spin Desalting Columns (89882, ThermoFisher), once for antibodies and twice for oligonucleotides. Activated antibodies and oligonucleotides were mixed and left to incubate overnight at 4°C.

### Comparison of Amplification Schemes

HT29 and SW403 EVs were captured on microarrays of CD63, CD9 (312102, Biolegend), goat anti-rabbit, EpCAM (MAB9601, R&D Systems) and CD81 (349501, Biolegend) antibodies and detected with CD9-BC51 antibodies (2 μg/mL) according to the EVPio workflow. Barcoded goat anti-mouse antibodies (GAM-BC23) were also used as a positive detection control. Amplification constructs were diluted down to 50 nM for labeling. Post-labeling, slides were washed for 3 min in a bath of washing buffer, dried, and imaged.

### Triplexed EVPio Assay

HT29 and SW403 EVs were captured on microarrays of CD63, CD9, goat anti-rabbit, EpCAM and CD81 antibodies and detected with barcoded CD63 (BC38), CD9 (BC51) and HSP70 (BC23; Invitrogen, PA5-34772) at concentrations of 1 μg/mL, 1 μg/mL, and 3 μg/mL, respectively, both separately (singleplexed detection) and simultaneously (triplexed detection). 2-branch amplification constructs were used for labeling at a concentration of 10 nM.

### HT29 and SW403 EV Profiling

HT29 and SW403 EVs were captured on microarrays of CD63, CD9, GFP (MA5-15256, Invitrogen), EpCAM and CD81 antibodies. The following trios of barcoded antibodies were used for triplexed detection: CD63-BC38, ITG α2 (MAB1233, R&D Systems)-BC64 and HSP70-BC23; CD9-BC51, ITG α6 (MAB1350, R&D Systems)-BC09 and Alix (MA1-83977, Invitrogen)-BC60; CD81-BC50, ITG β1 (MAB17781, R&D Systems)-BC27, HSP90-BC60; CD82 (342102, Biolegend)-BC43, ITG β4 (MAB4060, R&D Systems)-BC79, claudin-2 (32-5600, ThermoFisher)-BC61. Barcode-matched 2-branch amplification trees conjugated with Alexa Fluor 488, 555 and 647 were used for labeling.

### Data Analysis

Data was extracted from the microarray images using Array-Pro® Analyzer software (MediaCybernetics®). Further analysis was carried out using a custom MATLAB® script. First, intensity values were background-corrected by subtracting the median of the local background from the average value of each spot, and signal-to-noise (SNR) values were obtained by further dividing by the standard deviation of the local background. Next, signal intensity and SNR values were obtained for each condition by computing the mean and standard error of all associated spots from the background-corrected values. Signal intensity values were further corrected by subtracting the value of the matched no-EV control condition where applicable. Signal intensity and SNR values were thresholded using the value of the associated negative capture control plus two standard deviations. Plots were generated with custom MATLAB script with the exception of violin plots.^70^

## Associated Content

### Supporting Information

The following file is available online.

Detailed EVPio optimization data, including additional AR optimization conditions, duplex data with and without oligonucleotide labels and amplification, as well as buffer and oligonucleotide labeling optimization; additional data obtained in SW403 EVs; complete array images; experimental layout for the paneling experiment; reproducibility assessment; effect of AR on assay signal; oligonucleotide sequences (PDF).

## Author Information

### Author Contributions

The manuscript was written through contributions of all authors. R.M. performed the experiments and analyses. M.L.S. helped plan and run AR optimization. P.D.M. designed the oligonucleotide barcodes and helped design the oligonucleotide trees for branched DNA amplification. L.O.F.A. performed sample purification and cell culture. All authors have given approval to the final version of the manuscript.

### Notes

The authors declare no competing financial interest.

## Acknowledgments

We would like to thank the following individuals for their expertise and assistance with experimental design, reagent preparation and study planning: L. Alexandre, A. Wallucks, A. Ng. We also thank A. Ng for his help proof-reading this manuscript. We are grateful to J. Rak for supplying the A431-CD63-GFP cell line used in this work.

We are thankful for the financial support from Genome Canada and Genome Québec (Disruptive Innovation in Genomics) and the Natural Science and Engineering Research Council of Canada (NSERC Discovery grant, RGPIN-2016-06723). R.M. acknowledges funding from the Fonds de Recherche du Québec – Nature et Technologies (FRQNT), NSERC, and McGill University. D. J. acknowledges support from a Canada Research Chair.

## Abbreviations

EV: extracellular vesicle
EVPio: analysis of extracellular vesicle inner and outer proteins
SNR: signal-to-noise ratio
CRC: colorectal cancer
AR: antigen retrieval
PBS: phosphate buffer saline
TBS: tris-buffer saline
SSS DNA: sheared salmon sperm DNA
DS: dextran sulfate
EpCAM: epithelial cellular adhesion molecule
ITG: integrin
EGFR: epidermal growth factor receptor
ESCRT: endosomal sorting complexes required for transport
ELISA: enzyme-linked immunosorbent assay
HSP: heat-shock protein
PCR: polymerase chain reaction
RCA: rolling circle amplification.

