## Supporting information for "Extracellular Vesicle Antibody Microarray for Multiplexed Inner and Outer Protein Analysis"

**Table S1** Fluorescent detection signals for inner protein HSP90 and outer protein EGFR for different AR conditions following permeabilization with 0.05% Triton-X

| | | | HSP90 | EGFR |
|---|---|---|---|---|
| UREA | 50°C | 1 MIN | 0 | 3,100 |
| | | 2 MIN | 1,500 | 4,400 |
| | 70 °C | 1 MIN | 1,100 | 1,900 |
| | | 2 MIN | 1,000 | 2,100 |
| | 90°C | 1 MIN | 800 | 4,000 |
| | | 2 MIN | 2,100 | 2,600 |
| TBS | 50°C | 1 MIN | 0 | 5,600 |
| | | 2 MIN | 0 | 9,200 |
| | 70 °C | 1 MIN | 1,200 | 4,600 |
| | | 2 MIN | 1,600 | 3,200 |
| | 90°C | 1 MIN | 1,900 | 3,300 |
| | | 2 MIN | 1,700 | 3,400 |
| PBS | 50°C | 1 MIN | 4,000 | 14,800 |
| | | 2 MIN | 2,800 | 14,600 |
| | 70 °C | 1 MIN | 4,000 | 15,000 |
| | | 2 MIN | 11,300 | 12,300 |
| | 90°C | 1 MIN | 7,100 | 9,800 |
| | | 2 MIN | 7,800 | 10,900 |
| CONTROL (NO AR) | | | 0 | 5,000 |

| RFU[1] |
|---|
| 50,000 |
| 0 |

[1]A heatmap is superposed on the results. Signal values are above the threshold of negative capture control + 2SD unless greyed.

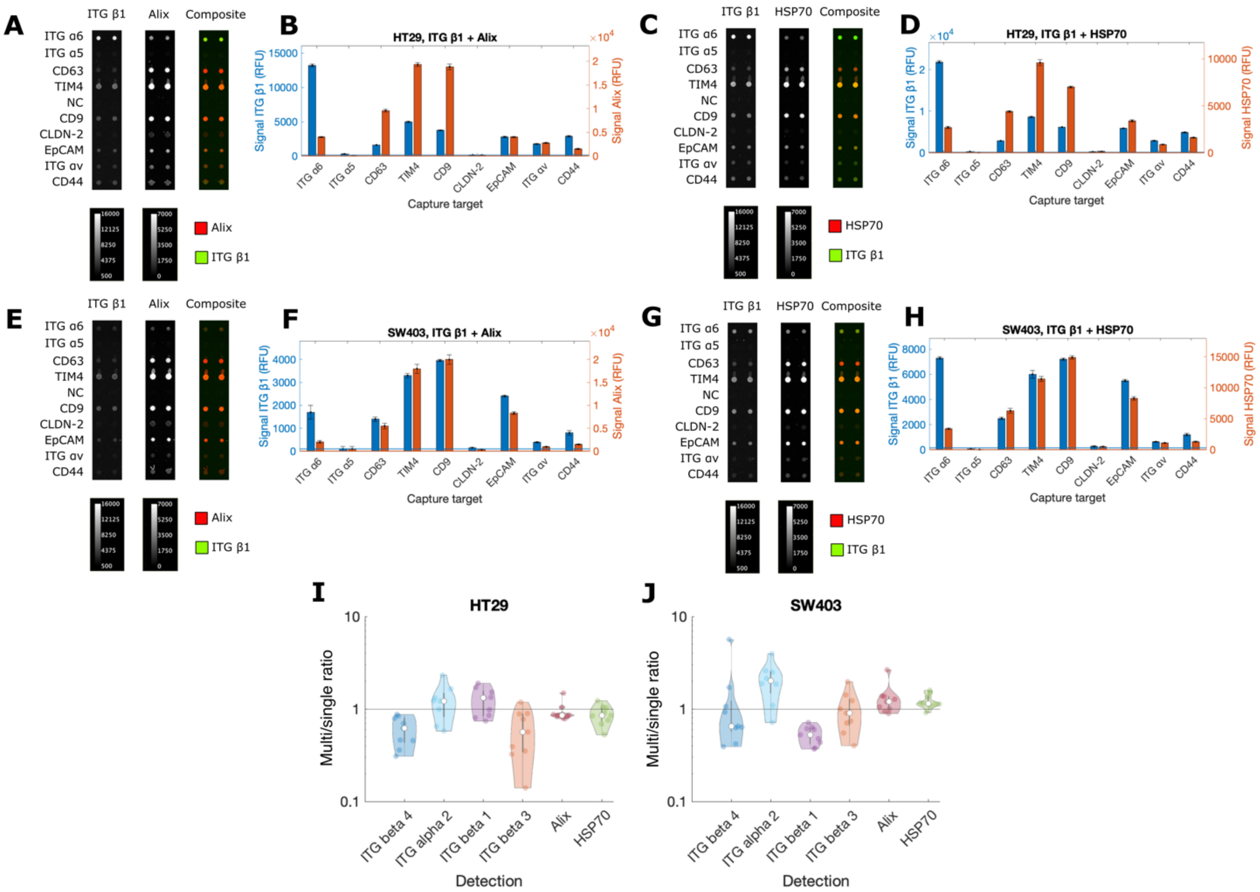

**Figure S1** EVPio analysis allows multiplexed detection of inner and outer EV proteins. (**A, C, E, G**) Micrographs and (**B, D, F, H**) quantification for the simultaneous detection of outer protein ITG β1 and inner proteins Alix or HSP70 for 9 capture antibodies on a microarray including a negative capture control (GFP) for (**A-D**) HT29 and (**E-H**) SW403 EVs. Inner proteins Alix and HSP70 were both successfully detected in combination with outer protein integrin β1. For both EV types, Alix and HSP70 were primarily detected in TIM4$^+$ and CD9$^+$ EVs. Integrin β1 signal was highest in integrin α6$^+$ HT29 EVs, and in integrin α6$^+$, TIM4$^+$ and CD9$^+$ SW403 EVs Thresholds for each detection target, calculated as the negative capture control (GFP) signal + 2SD, are indicated by horizontal lines color-matched to the corresponding bars. Error bars are SE. Ratios of the average multiplexed and singleplexed signals obtained for a same combination of detection antibody and amplification tree for all captured subpopulations of (**I**) HT29 and (**J**) SW403 EVs. Comparable singleplexed and multiplexed signals across detection targets and EV subpopulations emphasize the feasibility of multiplexed detection on EVPio. ITG, integrin.

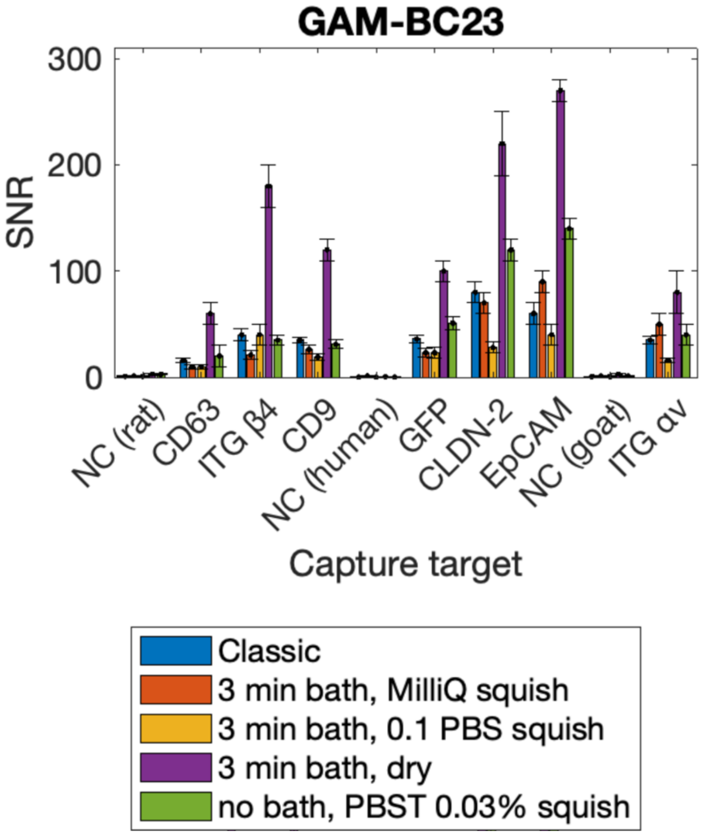

**Figure S2** Optimization of the final washing step for EVPio analysis for linear oligo labeling benchmarked for SNR using oligo-labeled goat anti-mouse (GAM) antibodies. Seven different mouse capture antibodies were microarrayed along with negative controls (rat, mouse and goat antibodies), and incubated with oligo-labeled GAM antibodies. Antibody binding was detected using a linear, fluorescently labelled oligo detection probe. 5 washing conditions were tested, including classic 15-min 0.03% Tween-20 PBS (PBST) bath wash followed by a quick MilliQ water rinse. A 3-minute wash in a bath of 0.03% PBST followed directly by a drying step yielded the highest overall SNR. NC, negative control. Error bars are SE.

**A**

| | Quenching | Incubation buffer | Hybridization buffer |
|---|---|---|---|
| Condition 1 | Normal | Normal | Normal |
| Condition 2 | + 0.5 mg/mL SSS DNA | + 0.5 mg/mL SSS DNA | + 0.5 mg/mL SSS DNA |
| Condition 3 | + 0.5 mg/mL SSS DNA | + 0.5 mg/mL SSS DNA | + 0.5 mg/mL SSS DNA + DS 0.05% |
| Condition 4 | Normal | Normal | 10 nM + DS 0.05% |
| Condition 5 | Normal | Normal | 0.27 nM + DS 0.05% |
| Condition 6 | + 0.5 mg/mL SSS DNA | + 0.5 mg/mL SSS DNA | 10 nM + DS 0.05% + 0.5 mg/mL SSS DNA |
| Condition 7 | + 0.5 mg/mL SSS DNA | + 0.5 mg/mL SSS DNA | 0.27 nM + DS 0.05% + 0.5 mg/mL SSS DNA |

**B**

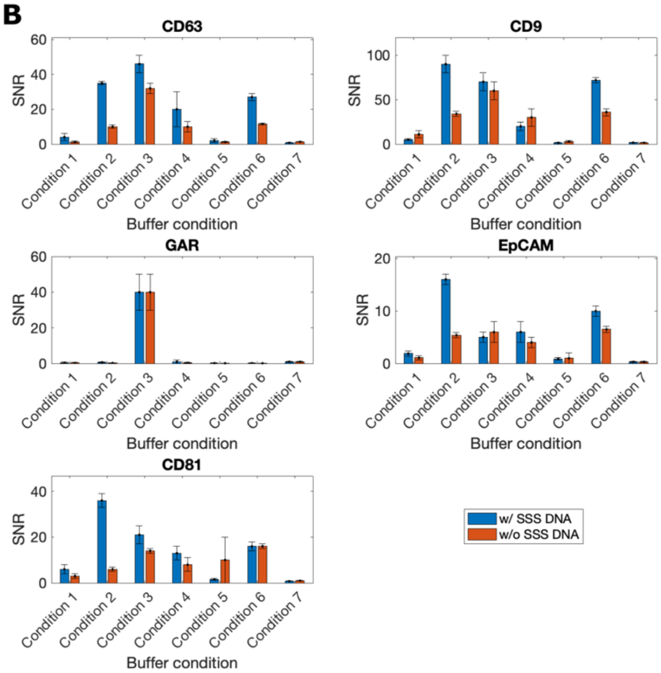

**C**

| | Quenching | Incubation buffer | Hybridization buffer |
|---|---|---|---|
| Condition 1 | + 0.5 mg/mL SSS DNA | + 0.5 mg/mL SSS DNA | + 0.5 mg/mL SSS DNA + DS 0.05% |
| Condition 2 | + 0.5 mg/mL SSS DNA | + 0.5 mg/mL SSS DNA | + DS 0.05% |
| Condition 3 | Normal | Normal | + DS 0.05% |
| Condition 4 | + 0.5 mg/mL SSS DNA | + 0.5 mg/mL SSS DNA | 10 nM + DS 0.05% + 0.5 mg/mL SSS DNA |
| Condition 5 | + 0.5 mg/mL SSS DNA | + 0.5 mg/mL SSS DNA | 10 nM + DS 0.05% |
| Condition 6 | Normal | Normal | 10 nM + DS 0.05% |
| Condition 7 | + 0.5 mg/mL SSS DNA | + 0.5 mg/mL SSS DNA | 2 nM + DS 0.05% |

**D**

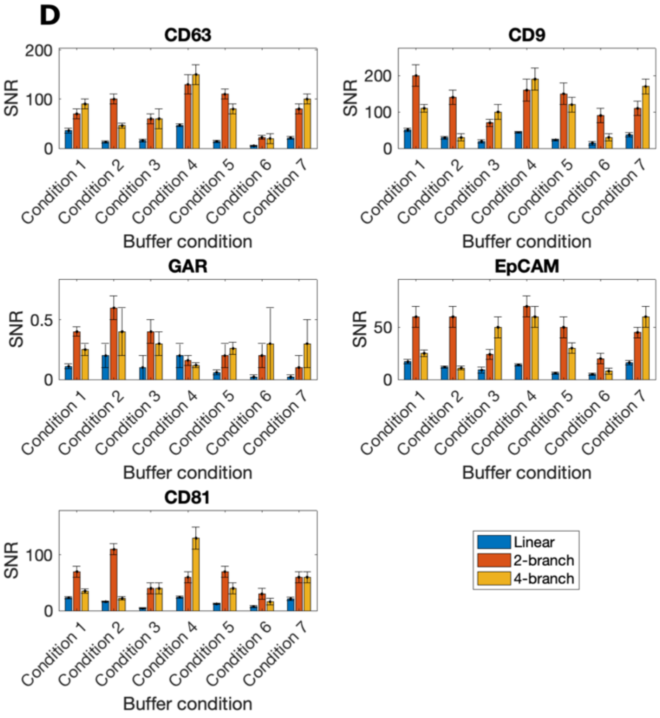

**Figure S3** Optimization of EVPio assay buffers towards maximizing SNR of linear, 2-branch and 4-branch oligo amplification using anti-CD63 detection antibodies. **(A-B)** Effect of the addition of sheared salmon sperm DNA (SSS DNA) and dextran sulfate (DS) to the buffers at the blocking (blue and orange bars), quenching, primary incubation and hybridization steps (numbered conditions) when the 2-branch tree is used for detection. Each bar graph is titled after one of the capture antibodies used, including goat anti-rabbit (GAR) as a negative capture control. Conditions 2 and 6 with SSS DNA at the blocking step led to the highest positive and lowest unspecific binding SNRs. **(C-D)** Comparison of the SNR of linear, 2-branch and 4-branch amplification constructs for different buffer compositions and/or fluorescent probe concentrations. Condition 4 (SSS DNA at all steps with 10 nM of fluorescent oligo construct (linear or tree) and DS in the hybridization buffer) yielded the highest overall detection SNRs with the lowest unspecific SNR. GAR is a negative control. Error bars are SE.

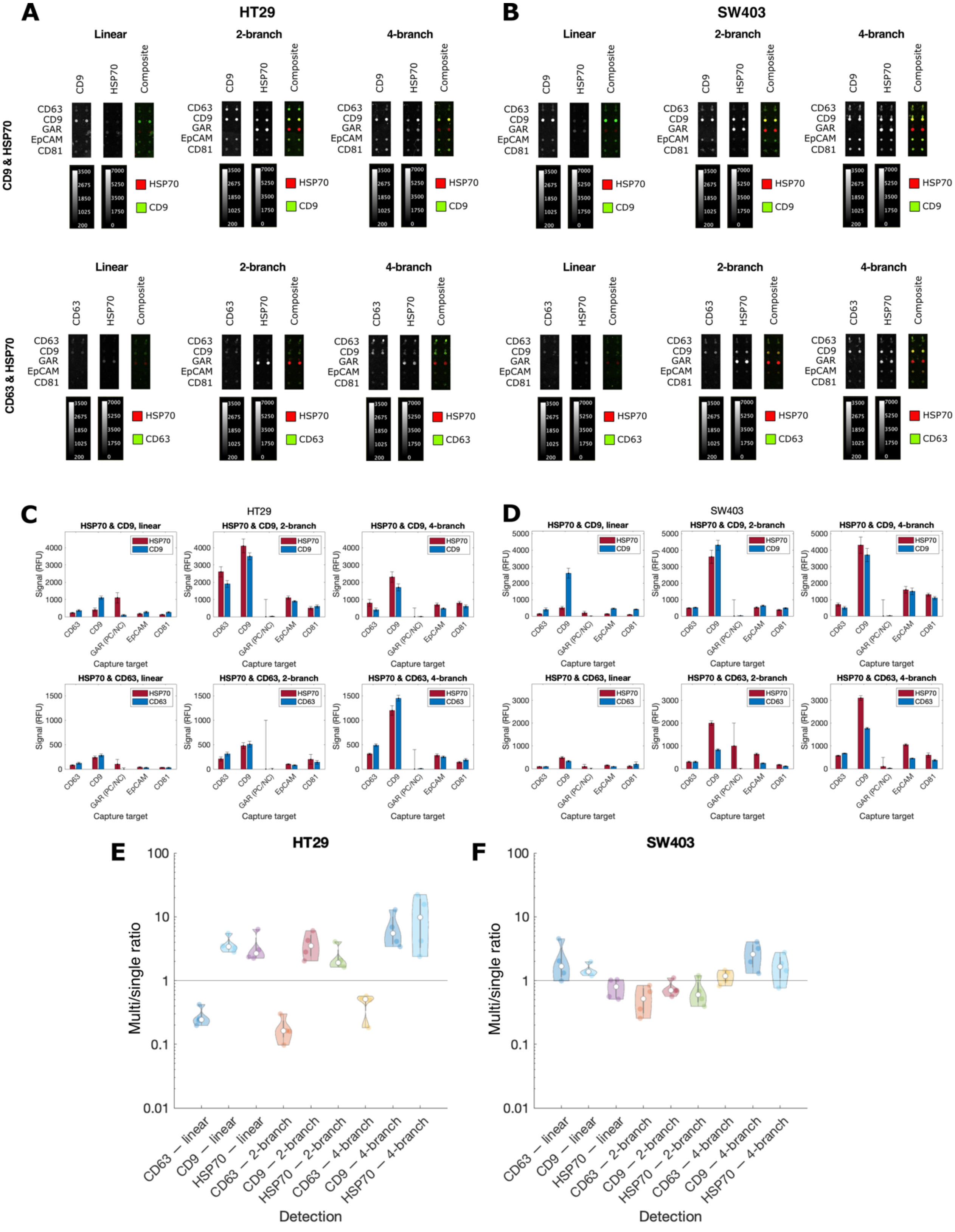

**Figure S4** EVPio analysis with duplexed, two-color detection. (**A-B**) Fluorescence micrographs and (**C-D**) quantification of detection in HT29 (*left*) and SW403 (*right*) EVs using (*left to right*) the linear, 2-branch and 4-branch oligo constructs for signal amplification. In micrographs, HSP70- and CD9-positive spots are colored red and green, respectively. Both targets were successfully detected simultaneously, with the 2-branch tree yielding consistently high signals. Ratios of the average multiplexed and singleplexed signals obtained for a same combination of detection antibody and amplification tree, averaged over all captured EV subpopulations, for (**E**) HT29 and (**F**) SW403 EVs. Comparable or enhanced signals were obtained for multiplexed conditions for most detection targets and EV subpopulations defined by capture antibodies, while some signal reduction was observed in a minority of cases. Some of the differences, positive or negative, may be due to experimental variation. Overall, results show the feasibility of multiplexed detection with branched DNA amplification for EVPio analysis. Error bars are SE.

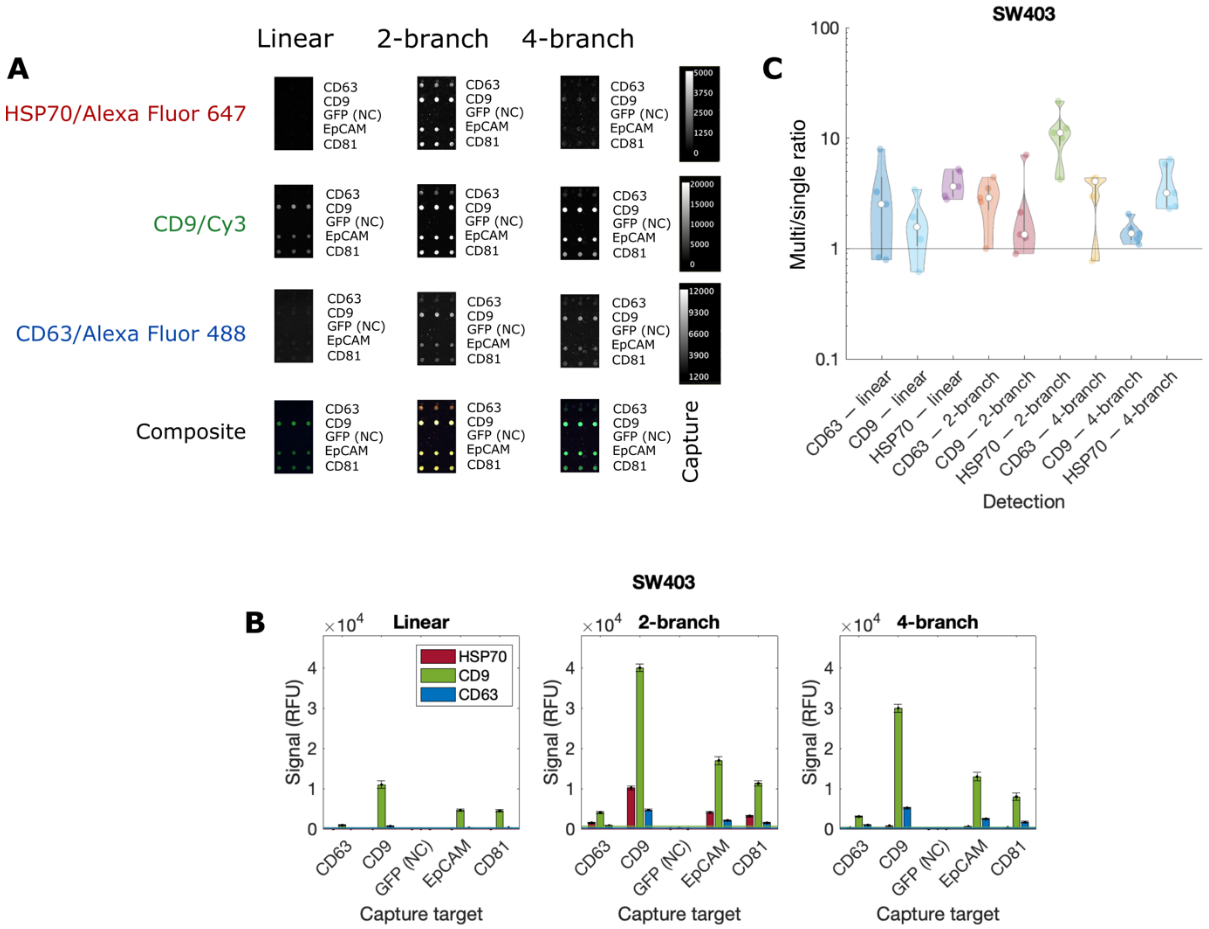

**Figure S5** Combinatorial EVPio analysis of inner, cytosolic proteins and outer, membrane proteins and validation of multiplexed versus singleplexed analysis. **(A)** Fluorescence micrographs of HSP70, CD9 and CD63 detected on EVPio microarrays with rows of CD63, CD9, CD81 and EpCAM capture spots, and negative controls (GFP) in SW403 EVs. Complete array images can be found in figure S6. **(B)** Bar graphs of (*left to right*) linear, 2-branch and 4-branch trees for signal amplification. 2-branch amplification gave overall higher signals. Errors bars are SE. **(C)** Violin log plots of the averaged (across capture targets) ratios of multiplexed over singleplexed signals for inner (HSP70) and outer (CD9, CD63) detection antibodies and different amplification trees. Averaged multiplexed signals are comparable, and up to ~ 10× higher than singleplexed signals.

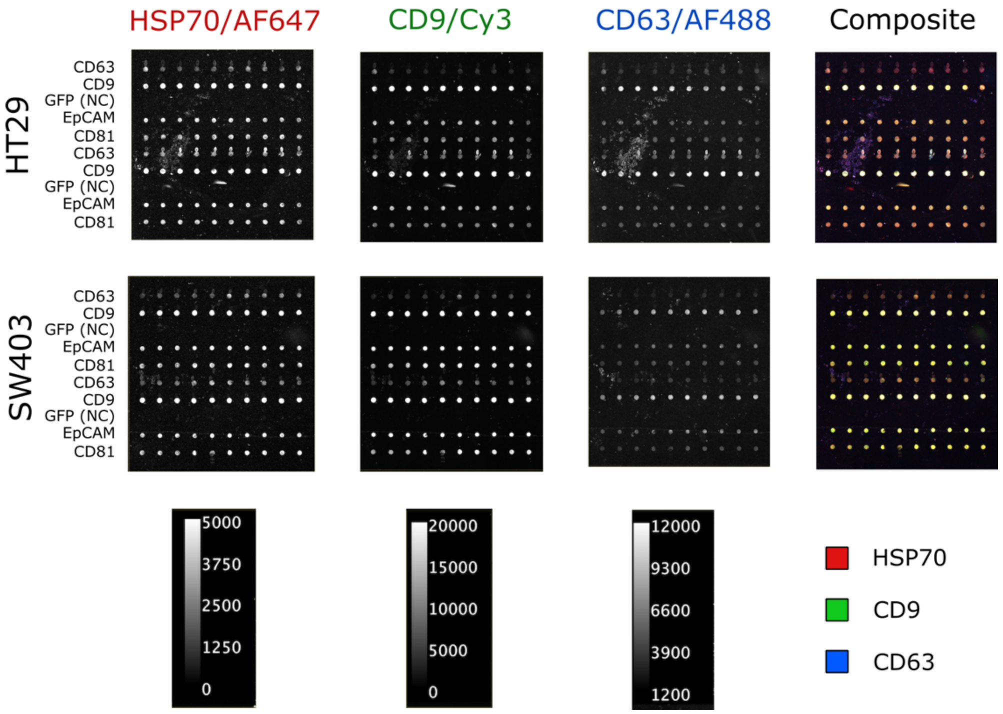

**Figure S6** Complete array micrographs for the triplexed EVPio analysis of HSP70, CD9 and CD63 in HT29 and SW403 EVs using the 2-branch oligo tree design.

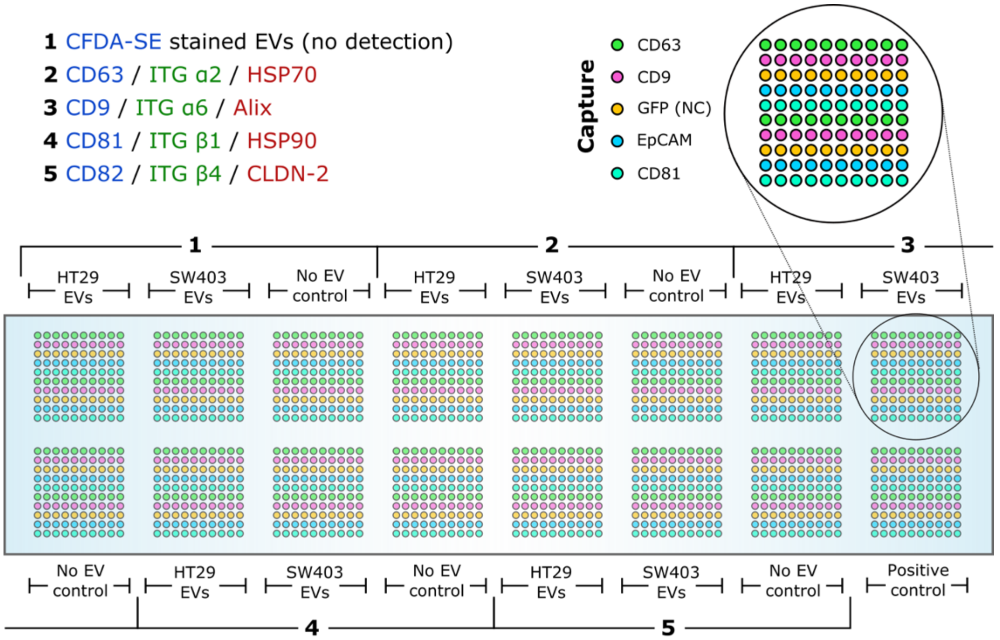

**Figure S7** Experimental EVPio analysis layout for HT29 and SW403 EV phenotyping. Each detection triplex (color-coded by fluorophore assignment; Alexa Fluor 488/CFDA-SE, Cy3, Alexa Fluor 647, *top left*) was assessed in a set of three wells, one per EV source (cell line) along with a negative detection control lacking EVs for signal correction. CFDA-SE is a non-specific label for EVs and was used in the first set of wells as a positive control to confirm EV binding. Capture antibodies were spotted on two lines of 10 replicate spot each (*top right*). A positive control well that is not incubated with sample, but with conjugated anti-mouse antibodies is used to monitor signal amplification (*bottom right*).

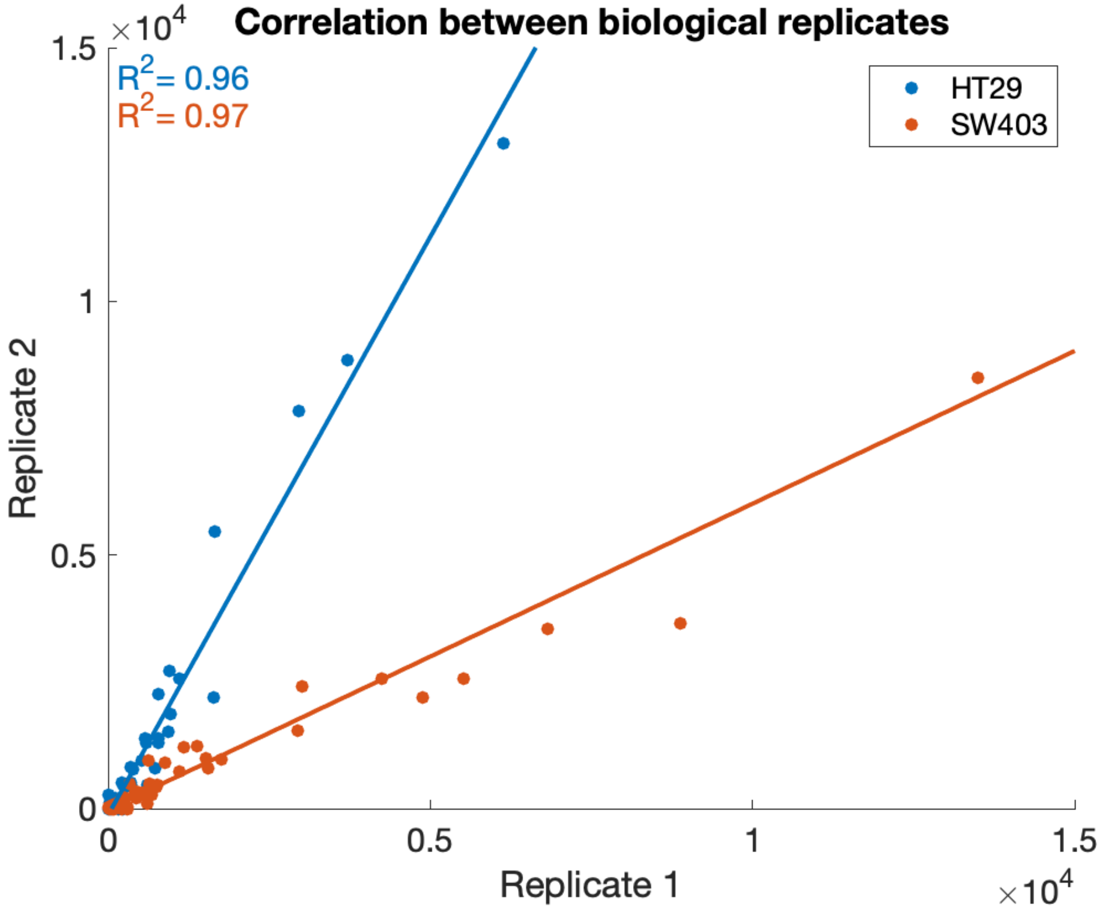

**Figure S8** Correlation between the two biological replicates of the EVPio panel for each type of cell line EV tested. Both HT29 and SW403 EVs had high coefficients of determination for the signals obtained in the two replicates, indicating good reproducibility.

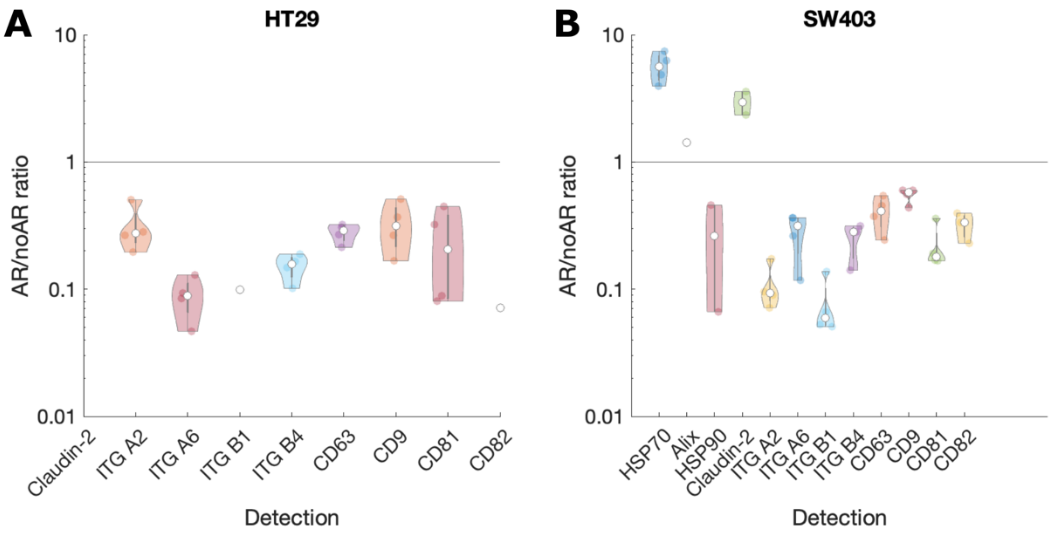

**Figure S9** Ratios of detection signals for EV outer proteins for assays with AR and without AR. The signal for outer proteins is significantly reduced in EVs from both cell lines. Inner targets tend to lead to AR/no AR ratios larger than 1, while outer target signals are most often reduced, hinting that AR is better used only on inner targets, and that assessing outer targets separately without AR can improve the performance of the assay.

**A**

| **Abbreviation** | **Name** | **Sequence (5' → 3')** |
|---|---|---|
| BC | Barcode | Variable |
| IBC | Intermediate barcode | CCTCCAAATAACCTTCTATCC |
| IBC1 | Intermediate barcode 1 | TGGTCTATTTATCCGCCGAAA |
| IBC2 | Intermediate barcode 2 | TACAATAGAACTGAGCGGAGA |
| FBC | Fluorophore barcode | TGGGAGGTTGGTGAGTGTGGA |
| TL | Toehold linker | GAGGAGAGT |
| ' | prime/reverse complement | – |

**B**

| **Strand name** | **Layout** | **Counts** |
|---|---|---|
| Imager | 5'-BC'-TL-FBC-3' | 1 |
| Fluorophore | fluorophore-5'-FBC'-3' | 1 |

**C**

| **Strand name** | **Layout** | **Counts** |
|---|---|---|
| Bridge | 5'-BC'-IBC-IBC-3' | 1 |
| Imager | 5'-FBC-FBC-T-IBC'-3' | 2 |
| Fluorophore | fluorophore-5'-FBC'-3' | 4 |

**D**

| **Strand name** | **Layout** | **Counts** |
|---|---|---|
| Bridge 1 | 5'-BC'-IBC1-IBC1-3' | 1 |
| Bridge 2 | 5'-IBC2-IBC2-IBC1'-3' | 2 |
| Imager | 5'-IBC2'-FBC-FBC-3' | 4 |
| Fluorophore | fluorophore-5'-FBC'-3' | 8 |

**E**

| **Barcode** | **Sequence (5' → 3')** |
|---|---|
| BC9 | ATGCTTCTACTCACT |
| BC23 | TAACACAAGCCAATG |
| BC27 | TTACGACCTACAATG |
| BC38 | TAGCCACTCATACGC |
| BC43 | TCCATACAACATCAG |
| BC50 | TCAGCACTTCCTTAC |
| BC51 | CGTCATTCACTACCT |
| BC60 | ATTATTCACGCACAC |
| BC61 | TAACATCCCTCATAG |
| BC64 | AATCATCCTATCTGC |
| BC79 | CGATTCCTAACTATT |

**Figure S10** Sequences of the oligonucleotides used for amplified detection. **(A)** Names and sequences of common segments**.** Composition of the **(B)** linear, **(C)** 2-branch and **(C)** 4-branch detection constructs. **(E)** Sequences of the 15-mer barcodes used for EVPio assays.
